# A bacteriophage cocktail significantly reduces *Listeria monocytogenes* without deleterious impact on the commensal gut microbiota under simulated gastro-intestinal conditions

**DOI:** 10.1101/2021.02.12.431056

**Authors:** Rasmus Riemer Jakobsen, Jimmy T. Trinh, Louise Bomholtz, Signe Kristine Brok-Lauridsen, Alexander Sulakvelidze, Dennis Sandris Nielsen

## Abstract

In this study, we examined the effect of a bacteriophage cocktail (tentatively designated FOP, for Foodborne Outbreak Pill) on the levels of *Listeria monocytogenes* in simulated small intestine, large intestine, and Caco-2 model systems. We found that FOP survival during simulated passage of the upper gastrointestinal was dependent on stomach pH, and that FOP robustly inhibited *L. monocytogenes* levels with effectiveness comparable to antibiotic treatment (ampicillin) under simulated ilium and colon conditions. FOP did not inhibit the commensal bacteria, whereas ampicillin treatment led to dysbiosis-like conditions. FOP was also more effective than antibiotic in protecting Caco-2 cells from adhesion and invasion by *L. monocytogenes*, while not triggering an inflammatory response. Our data suggest that FOP may provide a robust protection against *L. monocytogenes* should the bacterium enter the human gastrointestinal tract (e.g., by consumption of contaminated food), without deleterious impact on the commensal bacteria.

## Introduction

Antibiotics have been our main tool for the control of bacterial disease since their discovery in the late 1920s. However, their widespread use (and sometimes overuse) has also led to some major unintended consequences, such as increasing prevalence of resistant bacteria (1). In addition to the development of resistance, antibiotic treatment can heavily disrupt the ecology of the human microbiome by collateral damage to the commensal and symbiotic (2). The resulting dysbiosis might in turn increase the risk for developing diseases like obesity, asthma and inflammatory bowel disease (3–5). Consequently, there is an increasing interest in treatments that selectively target pathogenic bacteria, without disturbing the commensal gut microbiota of the gastrointestinal (GI) tract. Phage therapy is one such treatment option being explored, which involves the use of lytic bacteriophages to selectively kill disease-causing bacteria without impacting non-targeted benign bacteria that maybe beneficial for health (6, 7).

Bacteriophages (or “phages” for short) are viruses that attack bacteria in a host-specific manner, acting as self-replicating antimicrobials. Lytic phages replicate through the lytic cycle, where the phage infects the bacterial cell, uses the bacterial replication and translation machinery to replicate, and then lyses the cell to release new phage particles. Phages have the distinct advantage that they are (i) host specific, often only targeting specific strains within a specific species or (more seldom) within a limited number of related species and (ii) unable to infect and replicate in eukaryotic cells. These factors make phage therapy a promising means of targeted bacterial eradication within a microbial population without collateral damage to commensal bacteria (6). Furthermore, the mechanisms by which antibiotics and phages kill bacteria are fundamentally different, meaning potential bacterial resistances arise differently (8). Consequently, phages can kill bacteria that available antibiotics cannot, and the emergence of resistance to either would likely be mutually exclusive, allowing for the opportunity of phage-antibiotic complementary treatments (9). Phages may also be utilized for biocontrol applications, e.g., where phages are added to food products to reduce contamination with foodborne bacteria and consequent risk of foodborne diseases.

*Listeria monocytogenes* is a facultative gram-positive bacterium responsible for many cases of food-borne illness, manifesting as gastroenteritis, meningitis, encephalitis, mother-to-fetus infections, and septicemia. Although the annual number of *L. monocytogenes* infections globally is moderate, with 2,502 confirmed cases in the EU in 2017 (10) and an estimated 23,150 global cases in 2010, the mortality rate of infected individuals is considerable at 20-30% (11). The diverse clinical manifestations of *L. monocytogenes* are a result of its ability to enter both macrophages and other cell types, where it can survive and multiply (12). Crossing the epithelial barrier by adhering to and invading intestinal epithelial cells, gaining access to internal organs is the first step towards systemic infection of the host. In severe cases, the pathogen can also cross the blood-brain barrier to infect brain and meninges or cross the placental barrier to infect the fetus.

Preventive or therapeutic use of lytic phages is potentially an attractive approach for enhancing natural gut defenses against pathogenic bacteria such as *L. monocytogenes*, and/or as a complement to the current standard of care for various bacterial infections, including antibiotic treatment (13). For optimal efficacy, orally administered phages must first pass through several harsh environments during GI passage, including low pH in the stomach and pancreatic enzymes and bile salt in the small intestine. All these factors may reduce phage stability, destroying them or rendering them less active. Despite the long history of using phages therapeutically (14), the pharmacokinetics of orally administered phage preparations is still not well understood, and there is striking paucity of data on the impact GI passage on the phage viability and their ability to lyse their targeted bacteria in the GI tract after such passage.

The goals of this study were to test (i) survivability of phages under conditions mimicking those found in the stomach, (ii) potential of using the FOP bacteriophage cocktail to selectively target *L. monocytogenes* in the gut, using simulated human GI conditions (small and large intestines), and (iii) the ability of the same phage cocktail to protect Caco-2 cells from adhesion and invasion by *L. monocytogenes.*

## Materials and methods

### Bacteriophage cocktail

The FOP bacteriophage cocktail was created by Intralytix Inc. by combining, in approximately equal concentrations, three FDA-cleared commercial phage preparations currently marketed in the United States for food safety applications: ListShield™ (six phages active against *Listeria monocytogenes*), EcoShield PX™ (three phages active against Shiga toxin producing *E. coli* (STEC)), and SalmoFresh™ (six phages active against *Salmonella enterica*). Therefore, the FOP cocktail contains 15 unique lytic phages which together target *L. monocytogenes*, *Salmonella* spp., and STEC, including O157:H7 strains (**Table S1**). The FOP cocktail in liquid form is a clear to slightly milky in colour aqueous solution (pH 6.5-7.5) that was stored refrigerated (2-8°C) in the dark until use. For the stock FOP solution used in the experiments, the number of viable bacteriophages against *L. monocytogenes* was determined to be 10.83 PFU/ml by plaque assay (15) using *L. monocytogenes* strain LM114 on Luria-Bertani (LB)+ (10 g tryptone/L, 5 g yeast extract/L, 10 g NaCl/L, 0.02 M MgCl_2_, 0.001 M CaCl_2_, pH = 7.0) 1.5% agar.

### Bacterial strains

The *L. monocytogenes* strains, LM114 (serotype 4b) and LM396 (serotype 1/2a), were isolated from food processing plants, and provided by Intralytix. These strains were confirmed to be susceptible to the FOP cocktail via plaque assay (15). Both *L. monocytogenes* strains were propagated in LB broth (10 g tryptone, 5 g yeast extract, 10 g NaCl/L pH = 7.0) at 37°C with shaking (90 rpm). Quantification of all strains was performed via determination of colony forming units on LB agar.

### Consortium of small intestinal bacteria

To simulate a normal, healthy, small intestine microbiome, a consortium of 7 bacterial species were selected to represent a healthy ileal microbiota(16, 17) (**Table 1**). All bacteria were acquired from the German Collection of Microorganisms and Cell Cultures (DSMZ) and prepared and enumerated as described previously(18). 1 ml of consortium containing 10^8^ CFU ml^−1^ small intestinal bacteria was added to each TSI reactor.

**Table 1.** Small intestinal consortium bacterial strains, source, culturing time, and culture media.

| Species | Strain source | Origin | Culture time [h] | Culture media |
| --- | --- | --- | --- | --- |
| *Escherichia coli* | DSM 1058 | Human origin | 24 | GAM |
| *Streptococcus salivarius* | DSM 20560 | Blood | 6 | GAM |
| *Streptococcus luteinensis* | DSM 15350 | Human origin | 24 | GAM |
| *Enterococcus faecalis* | DSM 20478 | Human faeces | 24 | GAM |
| *Bacteroides fragilis* | DSM 2151 | Appendix abscess | 24 | GAM |
| *Veillonella parvula* | DSM 2008 | Human intestine | 48 | GAM |
| *Flavonifractor plautii* | DSM 6740 | Human faeces | 48 | GAM |

GAM = Gifu Anaerobic Medium. Modified from Cieplak et al. 2018 (18).

### Small intestinal model system

#### Small intestine in vitro simulation

To simulate passage of phages through the human stomach and small intestine, we used a recently developed dynamic *in vitro* model (TSI) (18), using fed state parameters (1h stomach passage, pH 4, bile salts = 10 mM, pancreatic juice = 100 U/ml) (18). For preliminary studies of phage viability during stomach passage, fed state parameters (30 minutes stomach passage at pH 2, bile salts = 4mM, pancreatic juice = 40 U/ml) were tested for comparison. The TSI model consists of five reactors with working volumes of 12 ml, each simulating the small intestine of one individual. pH and temperature were maintained at physiologically relevant levels, while simulated intestinal media, food, bile salts and digestive enzymes levels were established and maintained to simulate passage through stomach, duodenum, jejunum and ileum as previously described (18).

#### Bacteriophage impact on *L. monocytogenes* during stomach and small intestine passage

At the onset of simulated passage of the upper gastrointestinal tract, TSI reactors were inoculated with 0.5 ml of the FOP bacteriophage cocktail (10.81 log PFU/ml resulting in 10.03 log PFU/ml in the reactor) or ampicillin (500 mg/l in final solution), using saline solution (0.5 ml, 0.9% NaCl) as a control (**Figure 1A**). Before the ileal step 1 ml of *L. monocytogenes* strain LM396 suspension (7 log CFU/ml and 1ml of small intestinal consortium, was added into each reactor. Samples were taken from each reactor at the beginning and end of the ileum step and bacterial enumeration was performed by plate count on Palcam selective agar (19). The simulated small intestinal microbiota was enumerated using four different culturing media: Palcam Listeria Selective Agar (Palcam selective agar with Palcam selective supplement, Sigma-Aldrich) for enumeration of L. monocytogenes, Violet Red Bile Agar (VRB, Sigma Aldrich) for enumeration of E. coli, M17 Agar (M17, Oxoid) for enumeration of Streptococcus sp., MacConkey Agar (MCC, Sigma-Aldrich) for enumeration of E. faecalis, and Gifu Anaerobic Agar (GAM, NISSUI) where all species from the small intestinal consortium can be cultivated. Experiments were conducted for each preparation in triplicate.

**Figure 1.**
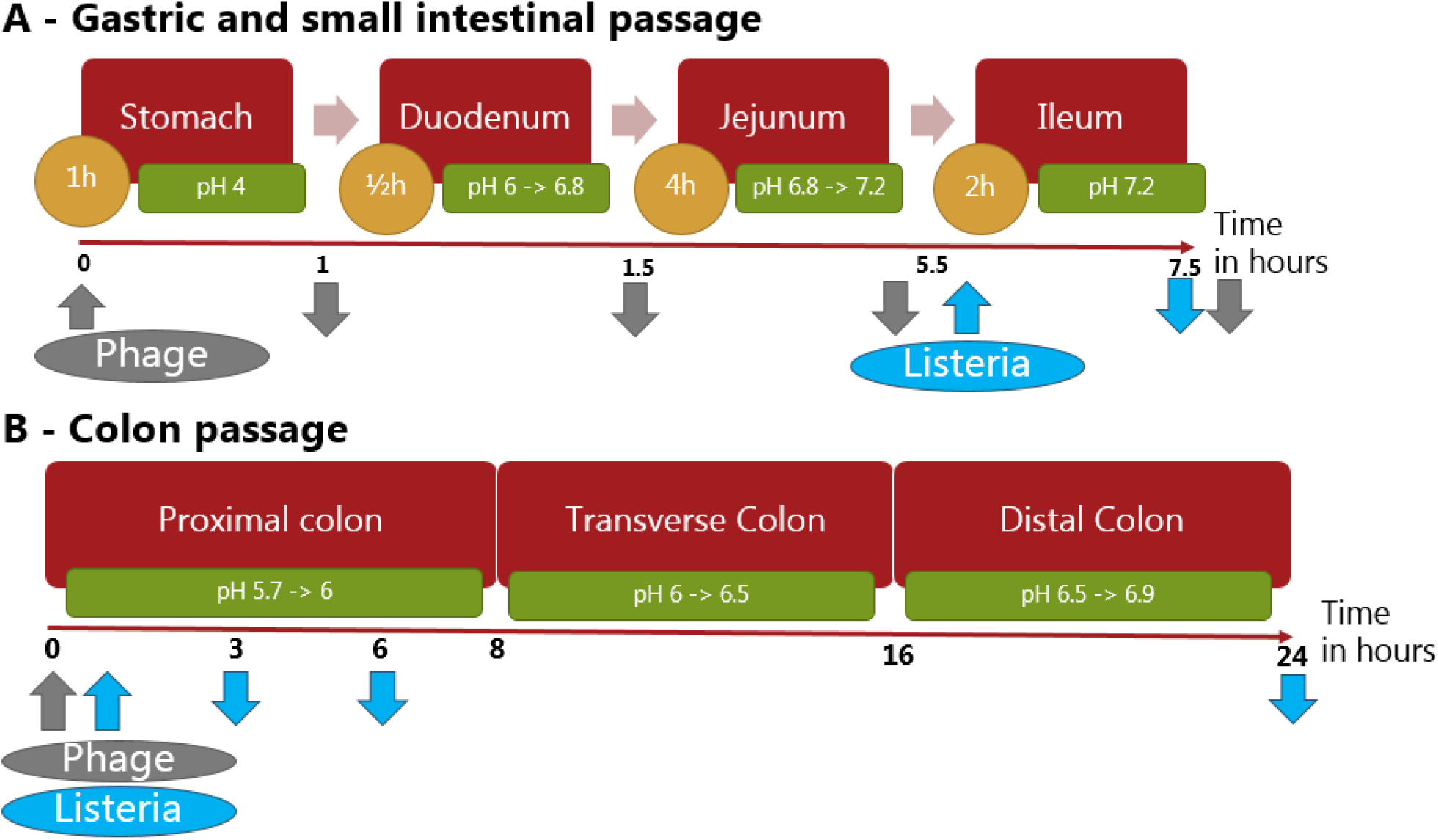
Flowchart of the (A) gastric and small intestine (TSI) and (B) colon (CoMiniGut) simulations showing *L. monocytogenes* (blue) and phage cocktail (grey) additions (upwards arrows) and sampling (downwards arrows) time points.

Preliminary experiments to test the persistence of the phage cocktail during gastric and small intestinal transit, and its efficacy under simulated intestinal conditions, were performed using only *L. monocytogenes* and phage cocktail, without adding the small intestinal bacterial consortium. 0.5 ml FOP was added at the beginning of the stomach stage, and samples were taken at the beginning of the jejunum, beginning of ileum, and at the end of the ileum stage (Figure 1A). Samples were diluted in SM buffer, refrigerated and the PFU was determined on the same day.

### Colon model system

#### Large intestine model system

To simulate colonic passage we used the CoMiniGut *in vitro* colon model (20). The CoMiniGut consists of five anaerobic reactors with working volume of 5 ml each. Reactors were filled with basal colon medium mixed with faecal inoculum from an anonymous adult donor (Ethical Committee (E) for the Capital Region of Denmark no. H-20028549) prepared as described (20) previously, and parameters like pH and temperature were monitored and maintained at physiologically relevant levels during simulations. In order to mimic the passage through the colon, pH was continuously controlled and gradually elevated from pH 5.7 to 6.9 over a 24 h period (20). Before the start of the experiment, the pH was adjusted to be 5.7 and anaerobic conditions were confirmed.

#### Impact of the FOP bacteriophage preparation on *L. monocytogenes* and colon microbiome

At the start of the experiments, combinations of 0.2 ml of the FOP phage cocktail (10^10^ pFU/ml in the reactors), 10^8^ CFU/ml *L. monocytogenes* strain LM 396, 0.5 ug/ml ampicillin or 0.9% saline controls were added and the colon simulation was run as previously described (20). Samples were taken at 3, 6 and 24h (**Figure 1B**). The number of viable *L. monocytogenes* cells was determined by plate count on PALCAM *Listeria* selective agar. To assess the impact of treatments on overall microbial composition samples, total bacterial DNA was extracted and subjected to 16S rRNA gene amplicon sequencing as described below.

### Sequencing of bacterial community

#### Library Preparation and Sequencing

The bacterial community composition was determined by Illumina NextSeq-based high-throughput sequencing (HTS) of the 16S rRNA gene V3-region, according to Krych et al. (21). Briefly, the amplified fragments with adapters and tags were purified and normalized using custom made beads, pooled, and subjected to 150 bp pair-ended Illumina NextSeq (V3 region 16S rRNA) sequencing.

The raw dataset containing pair-end reads with corresponding quality scores were merged and trimmed with usearch (22), using the following settings: -fastq_minovlen 100, -fastq_maxee 2.0, -fastq_truncal 4, and -fastq_minlen of 130 bp. De-replicating, purging from chimeric reads, and constructing de-novo zero-radius Operational Taxonomic Units (zOTU) were conducted using the UNOISE pipeline (23) and taxonomically assigned with sintax (24) coupled to the EZtaxon (25) 16S rRNA gene reference database. Sequences are available at the European Nucleotide Archive (ENA) with accession number PRJEB42055, https://www.ebi.ac.uk/ena/browser/view/PRJEB42055.

#### Bioinformatic Analysis

Initially the dataset was purged for zOTU’s, which were detected in less than 5% of the samples, but the resulting dataset still maintained 98 % of the total reads. Cumulative sum scaling (CSS) (26) was applied for the analysis of beta-diversity to counteract that a few zOTU’s represented a majority of count values, since CSS have been benchmarked with a high accuracy for the applied metrics (27). CSS normalisation was performed using the R software using the metagenomeSeq package (28). Alpha-diversity analysis was based on raw read counts, rarified to a median depth of 44574. R version 4.01 (29) was used for subsequent analysis and presentation of data. The data and code used is uploaded as supplementary data. The main packages used were phyloseq (30), vegan (31), ggpubr (32) and ggplot2 (33). Beta-diversity was represented by Bray Curtis dissimilarity.

### Caco-2 intestinal epithelial model

#### Caco-2 cell culturing

The human colon adenocarcinoma cell line Caco-2 (ATTC HTB-37, LGC standards, Middlesex, UK) at passage 53 was grown in Dulbecco’s Modified Eagles Medium (DMEM) supplemented with 10% (v/v) heat inactivated foetal bovine serum (FBS; Lonza, Basel, Switzerland), 1× Non Essential Amino Acids (NEAA), and 0.1 mg/mL gentamicin. Media was changed 3 times weekly. All solutions were obtained from Invitrogen, Gibco (Naerum, Denmark). The cells were cultured at 37 °C in a humidified atmosphere of 5% CO2.

#### Adhesion and invasion assay

Approximately 10^5^ Caco-2 cells were seeded in a 24-well microtiter plate in wells coated with 0.1% gelatin and grown for 14 days in DMEM supplemented with 20% HI-FBS, 10 mM HEPES, and an antibiotic-antimycotic solution (Gibco, Thermo Fisher Scientific) at 37°C in a humidified atmosphere maintained at 5% CO2. The medium was changed every 2-3 days. *L. monocytogenes* strain LM 396 was resuspended at 10^8^ CFU/mL in a solution comprising 80% DMEM with 10 mM HEPES and 10% FBS with appropriate amounts of the phage cocktail to achieve a MOI of 10, 100, or 1,000. After 30 min of pre-incubation, the *L. monocytogenes*-phage mixture was added to wells containing the Caco-2 cells and incubated for 1 h. The medium was then removed and saved for plate counting and cytokine measurements. The Caco-2 cells were washed with Dulbecco’s PBS (DPBS; Sigma-Aldrich, St. Louis, MO). Cells were then either lysed or incubated using DMEM with 10 mM HEPES containing 50 μg/mL gentamicin for 1.5 h. Cells treated with gentamicin were then washed with DPBS and lysed. For selected dilutions of the lysed cells that did not receive gentamicin, the lysed cells that received the antibiotic, or the medium removed prior to the first wash step (for the wells used for both the adhesion and invasion assay and the invasion assay), 100 μl was spread on LB agar plates and incubated at 37°C. After 48 h, the number of *L. monocytogenes* colonies was determined. Each trial was performed in triplicate.

#### Trans-epithelial resistance (TER) assay

The protective effect of FOP on the epithelial barrier exposed to *L. monocytogenes* was evaluated by measurement of TER using the Millicell Electrical Resistance System (Millipore, Bedford, MA) as previously described (34). To obtain polarized monolayers, Caco-2 cells were seeded onto Transwell filter inserts (0.4 μm pore size, 12 mm inside diameter, polycarbonate; Corning Incorporated, Corning, NY) at a concentration of 2 × 10^5^ cells/ml and cultivated for 14 days, with media change every 2 days. At 90-95% confluence, cells were moved into a cellZcope 2 next generation impedance-based cell monitoring unit. Treatments were performed after 2-3 days, once TER reached >1800 Ohm/cm^2^. Overnight cultures of bacteria were suspended in cell growth medium without antibiotics. A *L. monocytogenes* LM396 suspension in DMEM was added to the apical compartment at 10^6^ CFU/ml and incubated in a Forma Series 2 Water-Jacketed CO_2_ Incubator (Thermo Fischer, Waltham, MA) at 37 °C in a humidified atmosphere of 5% CO_2_. TER was measured before the addition of the bacteria (time zero) and then at 30 min time intervals and expressed as the ratio of TER at time *t* in relation to the initial value (at time zero) for each series. The net value of the TER was corrected for background resistance by subtracting the contribution of cell free filter and the medium (150 Ω). The TER of monolayers without added bacteria represented the control for each experiment. Experiments were performed with triplicate determinations.

### Statistics

R version 4.01 (29) was used for statitics and presentation of data (the data and code used is uploaded as supplementary material) using the vegan (31), ggpubr (32) and ggplot2 (33) packages. Analysis of variance (ANOVA) and (permutational ANOVA) permanova was used to evaluate group comparisons using Tukey’s range test and the Bonferroni–Holm method respectively for multiple testing correction. Significance was determined at P<0.05 level.

## Results

### The FOP bacteriophage cocktail selectively reduces *Listeria monocytogenes* in a small intestine *in vitro* model

The TSI (The Smallest Intestine) model was used to investigate the ability of the bacteriophage cocktail to endure digestive tract conditions and reduce *L. monocytogenes* levels in the ileum. The FOP phage cocktail was added before stomach passage, and *L. monocytogenes* was added at the beginning of the Ileum phase of the simulated small intestinal passage (**Figure 2A**). The bacteriophage cocktail caused a significant 1.5 log reduction in *L. monocytogenes* levels (q = 0.01) after two hours of ileal passage, while other representative ileal bacteria were not significantly affected (**Figure 2B**). Ampicillin treatment showed a similar 1.5 log reduction of *L. monocytogenes* (q = 0.01), but in contrast to the phage treatment, representative ileal bacteria also showed a 1.5 log fold reduction on average (**Figure 2B**). The small intestinal simulation was run using “fed” small intestine conditions (i.e., added food components, stomach pH 4, bile salts = 4mM; pancreatic juice = 40 U/ml), as stomach pH values of below 3.5 resulted in total phage deactivation (Figure S1). These “fed state” conditions use adjusted gastric pH values and bile salt concentrations to mimic GI conditions after a meal (18).

**Figure 2.**
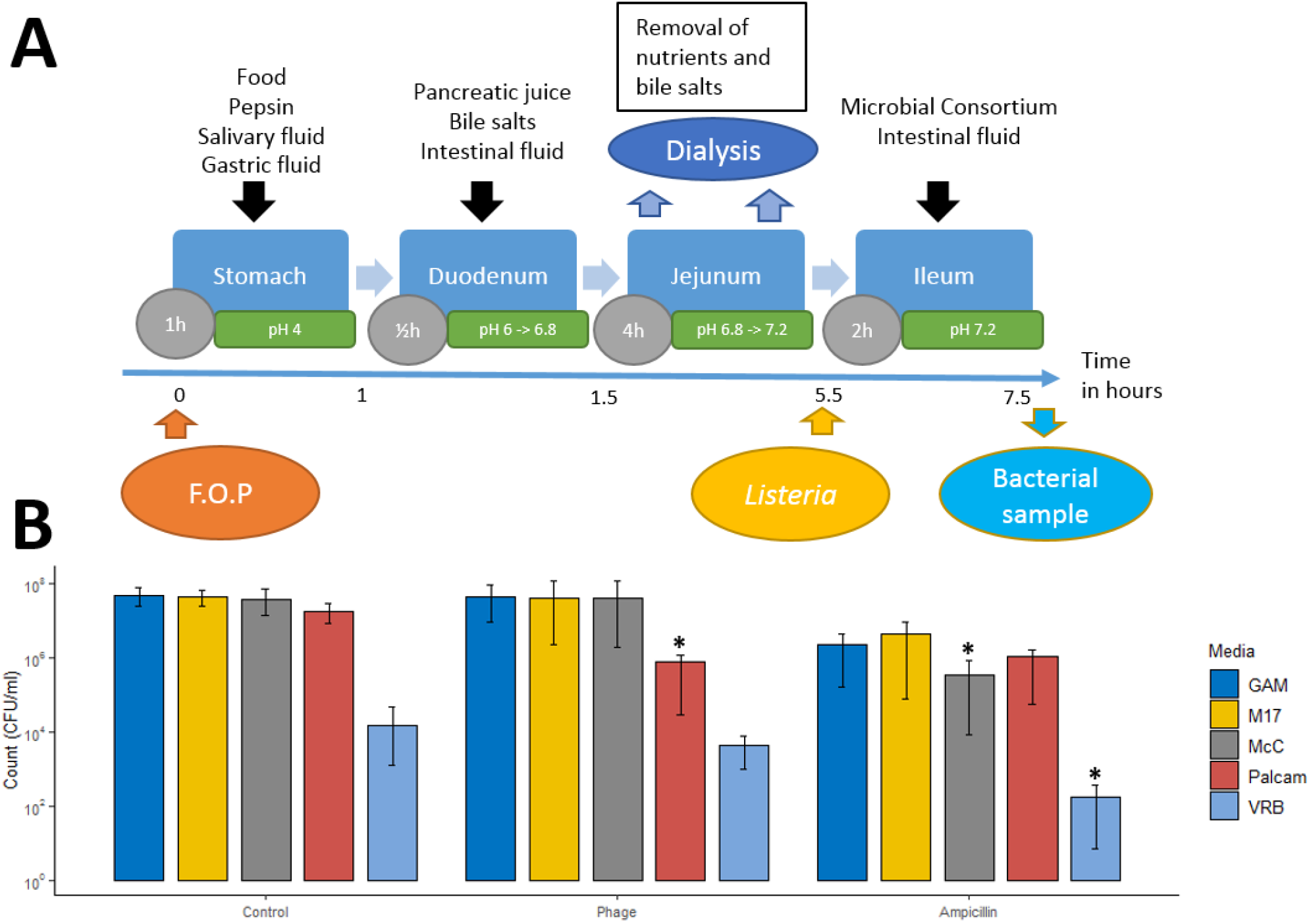
Impact of the FOP bacteriophage cocktail on *L. monocytogenes* in the ileum under simulated small intestinal conditions. A) Overview of experimental setup and sampling. B) Impact of the FOP and ampicillin treatment on *L. monocytogenes* and 7 representative bacterial species (Table 1) (6). The simulated small intestinal microbiota was enumerated using four different culturing media: Palcam *Listeria* Selective Agar (Palcam) for enumeration of *L. monocytogenes*, Violet Red Bile Agar (VRB) for enumeration of *E. coli*, M17 Agar (M17) for enumeration of *Streptococcus* sp., MacConkey Agar (MCC) for enumeration of *E. faecalis*, and Gifu Anaerobic Agar (GAM) where all species from the small intestinal consortium can be cultivated. All experiments were performed in triplicate. Significance calculated using one-way ANOVA using Tukey’s range test using non-treated samples with *L. monocytogenes* added as controls. *q< 0.05.

### The bacteriophage cocktail significantly reduces *L. monocytogenes* in a colon model while preserving bacterial community structure

To test the effect of the FOP phage cocktail on *L. monocytogenes* and the overall bacterial community in the colon, we used the CoMiniGut colon *in vitro* model. Phage cocktail, ampicillin or saline control was added at the start of the experiment, and the model was run for 24 hours to simulate colon passage. Samples treated with the bacteriophage cocktail had a 3-log reduction (p<0.01) in *L. monocytogenes* CFU at 3 hours, and a drastic 5-log reduction (p<0.01) after 24 hours of simulated colon passage (**Figure 3**). Ampicillin treatment resulted in a similar 2-log reduction of *L. monocytogenes* at 3 hours (p<0.01), with a final 5-log reduction (p<0.01) at 24 hours, compared to saline treated control samples.

**Figure 3:**
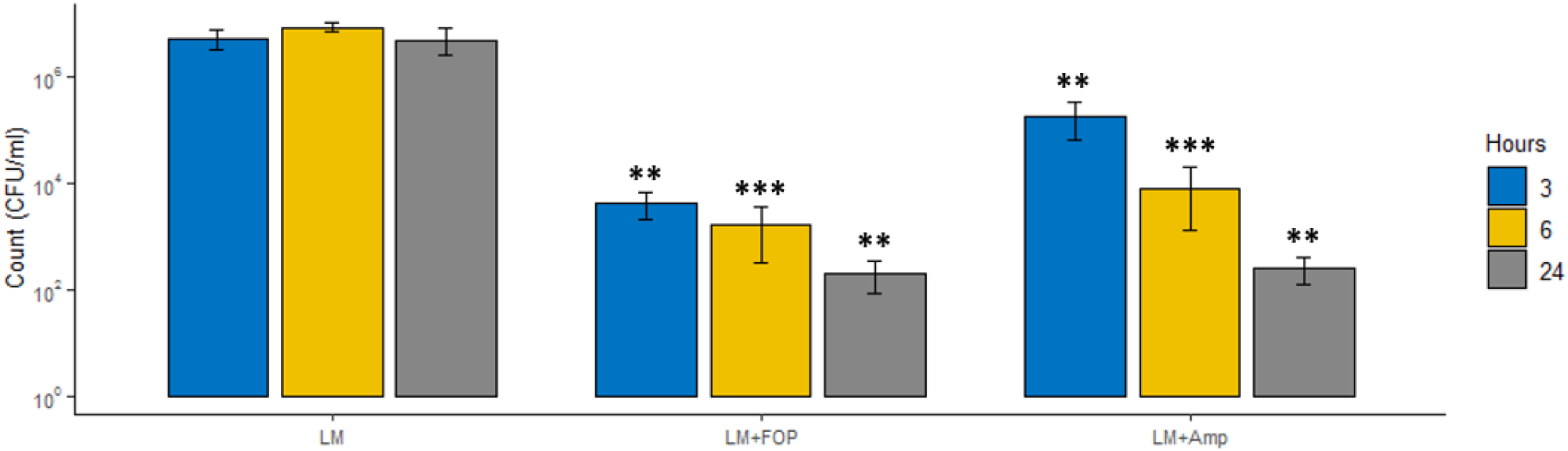
Impact of the FOP bacteriophage cocktail and ampicillin treatment on *L. monocytogenes* in the CoMiniGut *in vitro* colon model. Bacteria and treatments were added to the CoMiniGut reactors followed by sampling at 3, 6 and 24 hours. Experiments were performed in triplicate, and *L. monocytogenes* was enumerated by plate count on Palcam *Listeria* Selective agar. Significance calculated using one-way ANOVA using Tukey’s range test, using samples with *L. monocytogenes* added without treatment as control values. *q<0.05, **q<0.01, ***q<0.001.

To determine the overall impact of the phage cocktail on the colon bacterial community structure, 16S rRNA gene amplicon sequencing was performed. While there were shifts in bacterial community composition over time, *L. monocytogenes* was able to persist at relative abundance of approximately 25% throughout the 24 hours of simulated colon passage in non-treated (control) samples **(Figure 4A, B)**. The measured decrease in the relative abundance *L. monocytogenes* showed a similar trend to plate count results, with a decrease to 5% relative abundance at 6 hours and below detection limit (0.1%) at 24 hours **(Fig 4A, B)**. No significant effects of treatments were seen on alpha diversity, but overall, 24-hour samples showed a decrease in alpha diversity **(Fig S2)**. Noteworthy, at 24 hours the bacterial communities treated with the phage cocktail had community structure close to that of the untreated control (**Figure 4C**), while those treated with ampicillin markedly differed from on-treated controls (p = 0.002) (**Figure 4D**). Addition of *L. monocytogenes* had no significant effect on the overall bacterial community composition in non-treated samples at 24 hours (p = 0.24).

**Figure 4:**
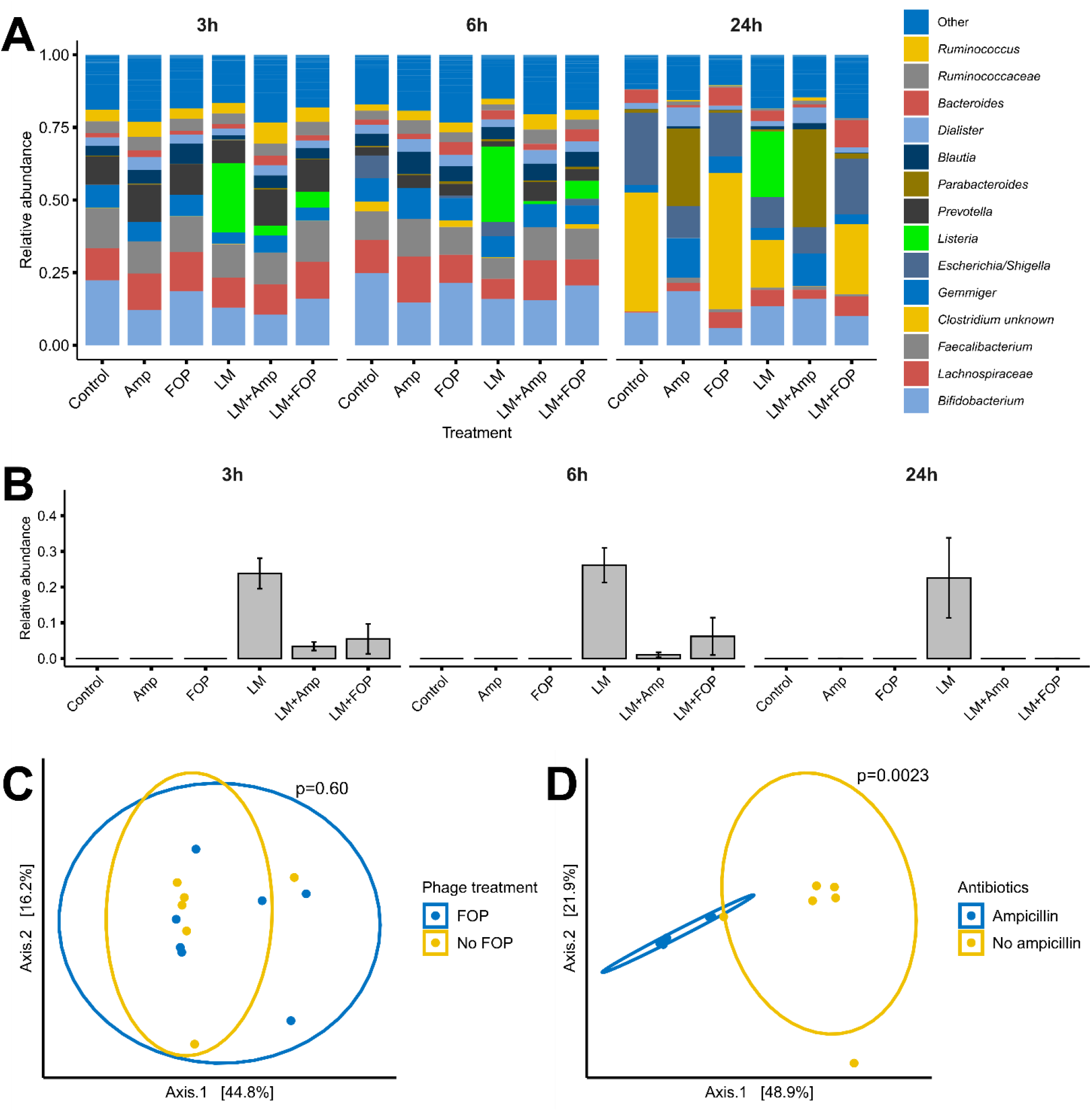
Impact of the FOP bacteriophage cocktail and ampicillin on the bacterial community in the CoMiniGut *in vitro* colon model, determined by 16S rRNA amplicon sequencing. Bacteria and treatments were added to the CoMiniGut reactors followed by sampling at 3, 6 and 24 hours. A) Relative abundance of bacterial genera ordered by average abundance. Legend colour order corresponds to chart order. B) Relative abundance of *L. monocytogenes* by treatment over 24 hours. C and D) PCoA plot of Bray-curtis dissimilarity metrics after 24h *in vitro* simulated colon passage with FOP or ampicillin treatment. Experiments were performed in triplicate and significance calculated using one-way ANOVA using Tukey’s range test.

### The FOP bacteriophage cocktail significantly reduces *L. monocytogenes* adhesion and invasion of Caco-2 cells

To examine if the FOP phage preparation may have protective effects in the intestine, we used Caco-2 epithelial cell line-based adhesion and invasion assays. Phage cocktail, ampicillin or PBS control was added to DMEM media containing *L. monocytogenes* and pre-incubated for 30 minutes, added to confluent Caco-2 cell monolayers and incubated for one hour. The bacteriophage cocktail resulted in a 5-log reduction (q<0.0001) in both adhesion and invasion of *L. monocytogenes*, while ampicillin treatment only resulted in a 1-log reduction (q<0.001) in adhesion and invasion (**Figure 5 A, B**). The reduction was highly dosage dependant with phage treatment at MOI 10 resulting in 1-log reduction (p>0.0001), while MOI 100 strongly reduced both adhesion and invasion (p>0.0001), and MOI 1000 prevented both adhesion and invasion (p>0.0001) (**Figure 5 C, D**).

**Figure 5:**
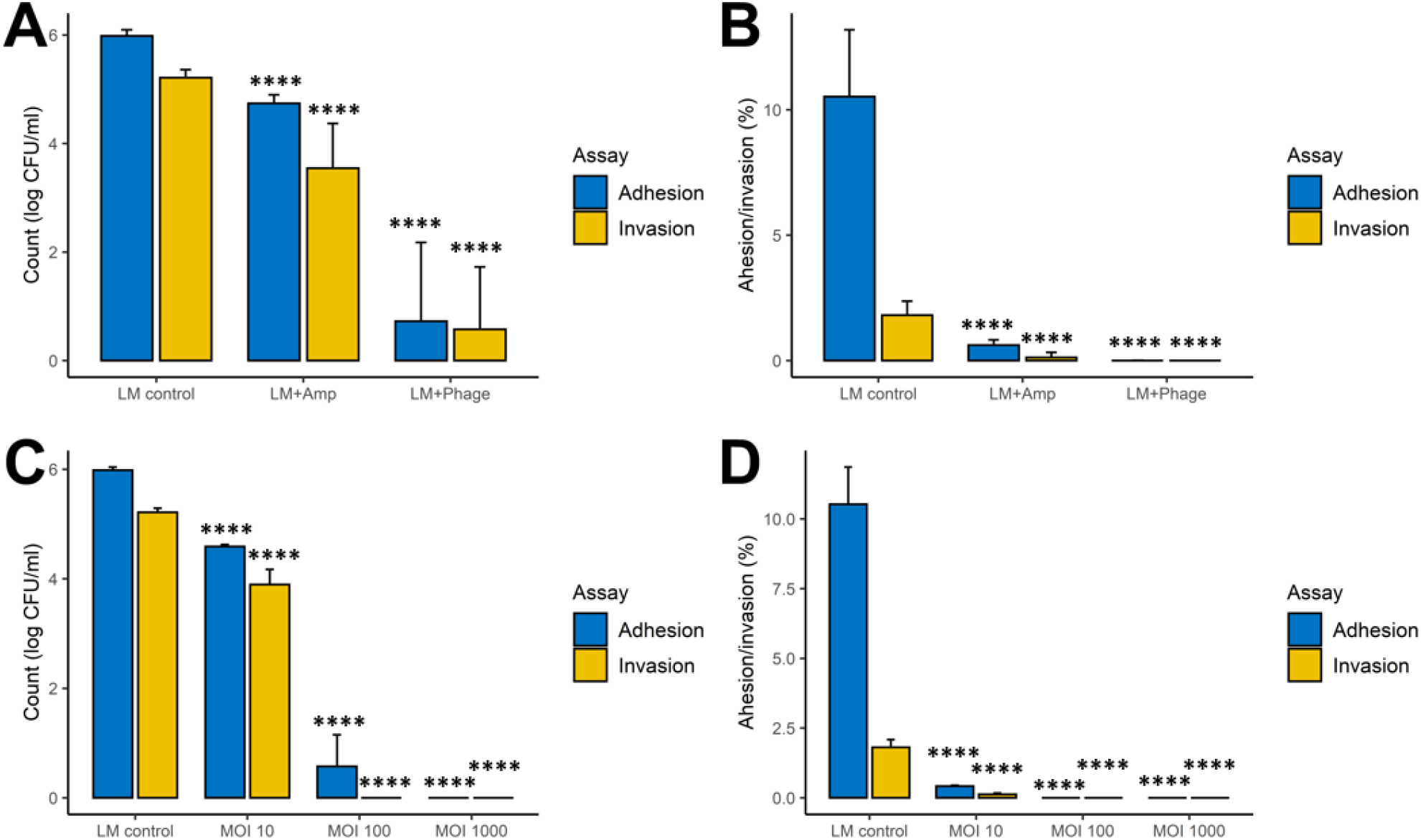
Impact of the FOP bacteriophage cocktail and ampicillin treatment on adhesion and invasion on a Caco-2 cell monolayer. Phage cocktail, ampicillin, or PBS control was added to a *L. monocytogenes* suspension in DMEM and pre-incubated for 30 minutes. The pre-incubated mixtures were added to wells and incubated for one hour. A) Adhesion and invasion CFU counts after 1 hour treatment with phage cocktail (MOI 100), ampicillin (500mg/L) or *L. monocytogenes* alone. B) Percentage adhesion and invasion after 1-hour treatment with phage cocktail (MOI 100), ampicillin (500mg/L) or *L. monocytogenes* alone. C) Adhesion and invasion CFU counts after 1 hour treatment with phage cocktail by MOI. D) Percentage adhesion and invasion CFU counts after 1-hour treatment with phage cocktail by MOI. Experiments were performed in triplicate, and *L. monocytogenes* was enumerated by plate count on Palcam *Listeria* Selective agar. Significance calculated using one-way ANOVA using Tukey’s range test, using samples with *L. monocytogenes* added without treatment as control values. *q< 0.05 – significant, **q<0.01, ***q<0.001.

### Cytokine production and trans-epithelial resistance in Caco-2 cells

To further assess the protective effect of the phage cocktail on the intestinal epithelium, we measured the cytokine response of Caco-2 cells after one hour of incubation with *L. monocytogenes* with or without 30 minutes of pre-incubation with the phage cocktail. However, we did not measure any cytokine response of neither IFN-γ, IL-1β, IL-6 nor TNF-α in regardless of treatment, with a detection limit of < 0.22 pg/ml (data not shown). Samples treated with the phage cocktail alone also had no detected effect on cytokine levels.

To measure the ability of the FOP phage cocktail in preserving epithelial integrity, we used a Caco-2 Transwell model to measure the effect on trans-epithelial resistance (TER) after exposure to *L. monocytogenes* with or without pre-incubation with the phage cocktail. Initial experiments showed that *L. monocytogenes* treatment led to a rapid drop in TER after 3 hours, but phage pre-treatment delayed this drop in a dosage dependent manner (**Figure S4A**). However, inspection of the wells after the experiment revealed an equally dosage dependent drop in pH. After the addition of HEPES buffer to stabilize pH, the drop in TER was delayed until after 24 hours (**Figure S4B**). pH measurements revealed that pH had decreased again at this time. We therefore concluded that *L. monocytogenes* was able to disrupt the integrity of the Caco-2 monolayer only by lowering the pH of media, and not through direct interaction with the cells. FOP appeared to have some dosage effect on preserving epithelial integrity, which was likely due to bacteriophages reducing the levels of and/or slowing the growth of the bacteria, delaying the lowering of pH.

## Discussion

In the present study we demonstrate that the FOP bacteriophage cocktail was able to survive gastric passage (under “fed” conditions) and selectively and significantly reduce *L. monocytogenes* in both the ileum and colon during *in vitro* simulated gastrointestinal tract passage. The bacteriophage treatment resulted in a significant reduction (up to 5 logs, p<0.01) of *L. monocytogenes* levels in both the ileum and colon model systems. Similar reductions were achieved by ampicillin treatment. However, the FOP phage cocktail specifically reduced only *L. monocytogenes* levels and had no impact on the commensal bacterial communities, whereas ampicillin indiscriminately affected both *L. monocytogenes* and the commensal bacterial communities.

In the TSI small intestinal *in vitro* model, treatment with the FOP cocktail led to a significant 1.5 log reduction in the *L. monocytogenes* levels during the relatively short 2-hour ileum transit time. In both the ileum and colon systems, the FOP cocktail only reduced *L. monocytogenes*, without an impact on any of the other commensal bacteria included in our system. In contrast, ampicillin treatment led to a significant killing-off of commensal small intestinal bacteria, in addition to a reduction of *L. monocytogenes*. In the CoMiniGut *in vitro* colon model, *L. monocytogenes* CFU counts were significantly reduced by 3 logs after 3 hours, and by 5 logs after 24 hours following bacteriophage treatment. Treatment by FOP did not alter the composition of the existing microbiome, demonstrating that FOP retains *L. monocytogenes*-specific bactericidal activity within the complex colonic bacterial community (in addition to *L. monocytogenes*, FOP also targets *Salmonella* spp. and STEC, but representatives of those pathogens were not included in our *in vitro* system). In the untreated samples, both CFU/ml and relative abundance of *L. monocytogenes* remained relatively stable over the 24-hour period. This is in contrast to other studies which have shown that *L. monocytogenes* is able to disrupt the composition of existing microbiota and infect robustly (35), and that certain probiotic intestinal species can inhibit the ability of *L. monocytogenes* to invade and grow (36). It is possible that we did not observe significant interaction between *L. monocytogenes* and colon bacteria because of the diluted conditions of the TSI and CoMiniGut models relative to the human GI tract.

Determination of the adhesion and invasion properties of *L. monocytogenes* to a Caco-2 epithelial cell monolayer showed that pre-treating *L. monocytogenes* with the FOP had a strong protective effect. This suggests that a sufficient intestinal bacteriophage concentration could prevent *L. monocytogenes* from invading the epithelial barrier. In support of this, a study using a mouse model with oral gavage with ListShield™ bacteriophage cocktail (the *L. monocytogenes*-targeting component of FOP) was able to reduce the concentration of *L. monocytogenes* in the GI tract, as well as its translocation to the spleen and liver (37). While high dosage / multiplicity of infection (MOI) appear to be vital for successful bacteriophage treatment (38), production of appropriately high titer preparations is feasible (39)(38).

We did not observe any significant effect of *L. monocytogenes* or the phage cocktail on neither trans-epithelial resistance (TER), nor cytokine production in the Caco-2 epithelial cell model. Previous studies have reported that *L. monocytogenes* strains producing listeriolysin O elicited a persistent IL-6 response (40). We do not know whether the LM396 strain we used during our studies produces listreiolisin O and/or its expression levels; thus, it is possible that the observed discrepancy between our study and the previous report (40) on the impact of *L. monocytogenes* on IL-6 production is due to LM396 not producing listeriolysin O or producing it in lower levels compared to the strains used in that previous study. With regards to the FOP phage preparation, our results support the idea that the phage cocktail itself does not provoke inflammatory response from the epithelial cells, as demonstrated by not eliciting IFN-γ, IL-1β, IL-6 nor TNF-α production in the Caco-2 cells.

Phage treatment is most commonly delivered orally, although many other modes of delivery such as auricular, intravesical, intrapulmonary, rectal, topical, and intravenous have also been used (41). We here show that the component of the bacteriophage cocktail targeting *L. monocytogenes* was able to survive gastric conditions, but only when stomach pH was 4 or above. The acidity of the human stomach is highly variable over time and between individuals, but stomach pH values of 4-5 are representative of conditions after meal ingestion (42). These results agree with the general observation that phage protein structures are acid labile (43, 44). Our data showed that the bacteriophages contained in FOP could survive “fed state” gastric conditions which simulate stomach and small intestinal conditions after a meal. However, phage titers declined more rapidly in the “unfed” gut conditions, suggesting that phage efficacy may be significantly reduced by more acidic stomach conditions. Furthermore, it is possible that some of the GI conditions not examined in our model systems such as intestinal peristalsis, complex microbiota, various diets, etc. may further reduce phage viability – and thus efficacy - *in vivo*. One approach commonly used during therapeutic phage applications in the former Soviet Union and Eastern Europe (and during some animal studies) was to administer oral phage preparations together or shortly after administering sodium bicarbonate to reduce stomach acidity. Alternatively, there is currently a wide range of encapsulation methods available that could be used to formulate lytic phage preparations in “enteric” gel caps or tablets, to ensure phage survival through the stomach and their release in the intestine (45).

In summary, our data demonstrate that the FOP bacteriophage cocktail is able to (i) endure gastric passage under “fed” conditions, and (ii) significantly reduce *L. monocytogenes* in a highly selective manner under *in vitro* human gastric conditions while having no detectable deleterious effect on the commensal gut microbiota. Furthermore, the data suggest that the phage cocktail has a strong protective effect on adhesion and invasion by *L. monocytogenes* through a Caco-2 monolayer. These results are in agreement with previous reports that FOP can provide robust protection against pathogenic bacteria both *in vivo* (46) and *in vitro* (47), while avoiding detrimental effects on the existing microbiota. Taken together, our data provide further support to the idea that lytic phages may provide some important health benefits, e.g., when consumed as dietary supplements, by enhancing natural defenses of the GI tract against specific foodborne bacterial pathogens. For example, phages with strong lytic potency against *L. monocytogenes* included in the FOP preparation may help increase gut resilience against *L. monocytogenes* by specifically killing these bacteria (and preventing their attachment and invasion into epithelial cells) if they are introduced into the gut via consumption of contaminated food. Such nutraceutical products (FOP or similar) may be taken routinely as a regular gut enhancing supplements, or during outbreaks of foodborne diseases or food recalls caused by bacterial pathogens targeted by the lytic phages in those supplements and may help reduce the risk of foodborne illness caused by major foodborne bacterial pathogens.

## Acknowledgements

AS holds an equity stake in Intralytix, Inc. JTT is an employee at Intralytix Inc. Funding for the study was provided by Intralytix, Inc. The phage preparations contained in the FOP are the subject of several issued and pending patent applications.

